# Defining Microbial Community Functions in Chronic Human Infection with Metatranscriptomics

**DOI:** 10.1101/2023.06.06.543868

**Authors:** Aanuoluwa E. Adekoya, Hoody A. Kargbo, Carolyn B. Ibberson

**Affiliations:** Department of Microbiology and Plant Biology, University of Oklahoma, Norman, OK 73019

**Keywords:** human infection, transcriptomics, chronic wounds, cystic fibrosis, microbial community functions

## Abstract

Chronic polymicrobial infections (cPMIs) harbor complex bacterial communities with diverse metabolic capacities, leading to competitive and cooperative interactions. Although the microbes present in cPMIs have been established through culture-dependent and -independent methods, the key functions that drive different cPMIs and the metabolic activities of these complex communities remain unknown. To address this knowledge gap, we analyzed 102 published metatranscriptomes collected from cystic fibrosis sputum (CF) and chronic wound infections (CW) to identify key bacterial members and functions in cPMIs. Community composition analysis identified a high prevalence of pathogens, particularly *Staphylococcus* and *Pseudomonas*, and anaerobic members of the microbiota, including *Porphyromonas, Anaerococcus, and Prevotella*. Functional profiling with HUMANn3 and SAMSA2 revealed that while functions involved in bacterial competition, oxidative stress response, and virulence were conserved across both chronic infection types, ≥40% of the functions were differentially expressed (padj < 0.05, fold-change >2). Higher expression of antibiotic resistance and biofilm functions were observed in CF, while tissue destructive enzymes and oxidative stress response functions were highly expressed in CW samples. Of note, strict anaerobes had negative correlations with traditional pathogens in both CW (*P* = -0.43) and CF (*P* = -0.27) samples and they significantly contributed to the expression of these functions. Additionally, we show microbial communities have unique expression patterns and distinct organisms fulfill the expression of key functions in each site, indicating the infection environment strongly influences bacterial physiology and that community structure influences function. Collectively, our findings indicate that community composition and function should guide treatment strategies for cPMIs.

**Importance:** The microbial diversity in polymicrobial infections (PMIs) allows for community members to establish interactions with one another which can result in enhanced disease outcomes such as increased antibiotic tolerance and chronicity. Chronic PMIs result in large burdens on health systems, as they affect a significant proportion of the population and are expensive and difficult to treat. However, investigations into physiology of microbial communities in actual human infection sites is lacking. Here, we highlight that the predominant functions in chronic PMIs differ, and anaerobes, often described as contaminants, may be significant in the progression of chronic infections. Determining the community structure and functions in PMIs is a critical step towards understanding the molecular mechanisms that drive microbe-microbe interactions in these environments.

---

Microbes live in multi-species communities where community structure and function dictate key processes such as nutrient cycling, tolerance to disturbances, and in infection sites, disease progression. The presence of diverse microbes with a wide range of metabolic capacities and large nutrient gradients often leads to microbe-microbe interactions in chronic polymicrobial infections (cPMIs) (1). These cooperative and competitive interactions can result in increased disease severity, increased antimicrobial tolerance, and chronicity compared to single species infections (2,3). Although we have known that chronic infections are composed of polymicrobial communities for over 100 years, pathogenesis research has focused on the physiology of a handful of well-known pathogens in isolation in laboratory and animal models, and data on microbial community physiology in human infection sites is lacking (4,5,6). Further, the contribution of the normal flora identified in cPMIs to disease progression has remained debatable, and members of the microbiota are often ignored in current treatment plans (7). Therefore, two important knowledge gaps are the key functions that drive each chronic PMI and the metabolic activities of the array of microbes present. To address these questions, we analyzed 102 previously published metatranscriptomes collected from people with CF (CF: 30%) and chronic wound infections (CW: 70%) to identify key bacterial members and community functions in these typical examples of clinically important cPMIs (8,9).

## Anaerobes are prominent in chronic infections

We identified microbial communities in 90 of our 102 samples (CF:31 CW:59) through community composition analysis with MetaPhlAn4 (Table S1). Identification of the genera present revealed that both the CW and CF sputum samples contained a mix of traditional pathogens from the genera *Staphylococcus, Pseudomonas*, and *Streptococcus*, along with anaerobic members of the microbiota (Fig. 1A), concordant to what is expected in these infections based on previous metagenomic and 16S rRNA gene data (8,10,11,12,13). While the mean number of species identified in each sample aligns with previous reports (10,12,13,14,15), we found the CF samples were more diverse than CW samples with a mean of 11.8 and 6.7 species identified, respectively (*P*-value < 0.01) (Fig. 1B). The increased diversity in CF sputum compared to CW wounds was also observed with both Shannon and Simpson diversity indices (Fig. 1C&D). Interestingly, we identified a high abundance of transcripts assigned to anaerobes in these samples (Fig 1A&E, Table S1), suggesting the chronic infection environments are likely hypoxic. Further, we found that while anaerobes co-occurred with traditional pathogens in over 50% of samples (CF: 80.7%, CW: 52.5%), there was a strong negative correlation between the anaerobes and traditional pathogens in both sites, indicating possible competitive interactions (Fig. S1).

**Figure 1:**
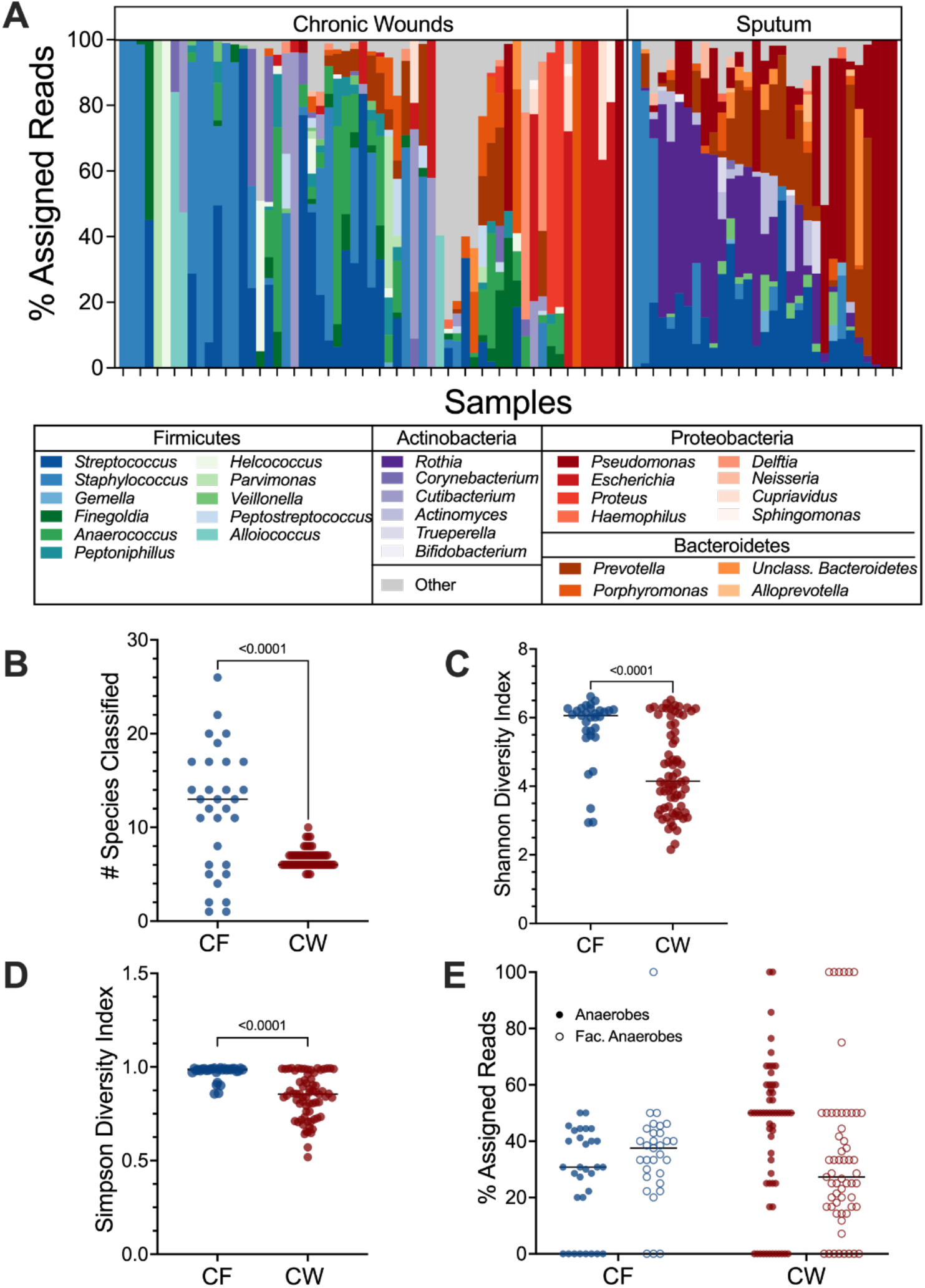
Bacterial community composition in CF and CW environments. **A)** Relative abundance of bacterial genera present in at least 3 samples with a % assigned read abundance of at least 1%. 29 genera were identified in CF samples and CW 36 in wound samples. **B)** Distribution of the number of species with a relative abundance of at least 1% in CF and CW samples. **C)** The Shannon diversity index of each sample. **D)** Distribution of the Simpson diversity index in each sample. **E)** Distribution of the percentage of reads assigned to anaerobes (closed circles) and facultative anaerobes (open circles) in each sample in the CF and CW environments. For plots B-E, CF samples are in blue and CW samples are in red. P-values and brackets indicate comparisons that were deemed statistically significant (T-test, *P*-value <0.05)

## CF sputum has increased expression of antibiotic resistance and biosynthetic pathways while tissue destructive and catabolic pathways are primarily expressed in CW infections

Through profiling with both SAMSA2 and HUMANn3, we classified the level 4 enzyme commission (EC) functions in each sample (SAMSA: 4527, HUMANn3: 2459). Our analysis revealed that several EC classes involved in oxidative stress responses, virulence, bacterial competition, fatty acid metabolism, and iron acquisition were conserved across infection environments (Table S2), indicating that bacterial community members in these infection types may be competing with one another for resources while tolerating host innate immune mechanisms and simultaneously expressing their virulence functions. However, while some key functions were conserved across both infection sites, over 40% of the functions identified were differentially expressed (qvalue < 0.05, fold-change > 2) between the two sites (40.4% and 43.0% for SAMSA2 and HUMANn3, respectively), with the majority displaying higher expression in the wounds compared to the sputum. There were key differences in the types of functions that were highly expressed in each site. CF sputum displayed high expression of antibiotic resistance functions, iron acquisition, virulence factors, and functions important for attachment to host surfaces (Fig. 2A & Table S2). In contrast, CW infections had high expression of functions involved in oxidative stress response and tissue destructive enzymes and virulence factors (Fig. 2A & Table S2). Taking a deeper look into the expression of metabolic pathways in each site revealed the enrichment of catabolic pathways, such as the glycogen degradation pathway and the valine degradation pathway, as well as catabolism support pathways, including phospholipase synthesis, in CW samples (Table S3). In contrast, in the CF samples there was an enrichment of biosynthetic pathways, such as the fatty acid elongation pathway, oleate, palmitoleate and valine biosynthesis.

**Figure 2:**
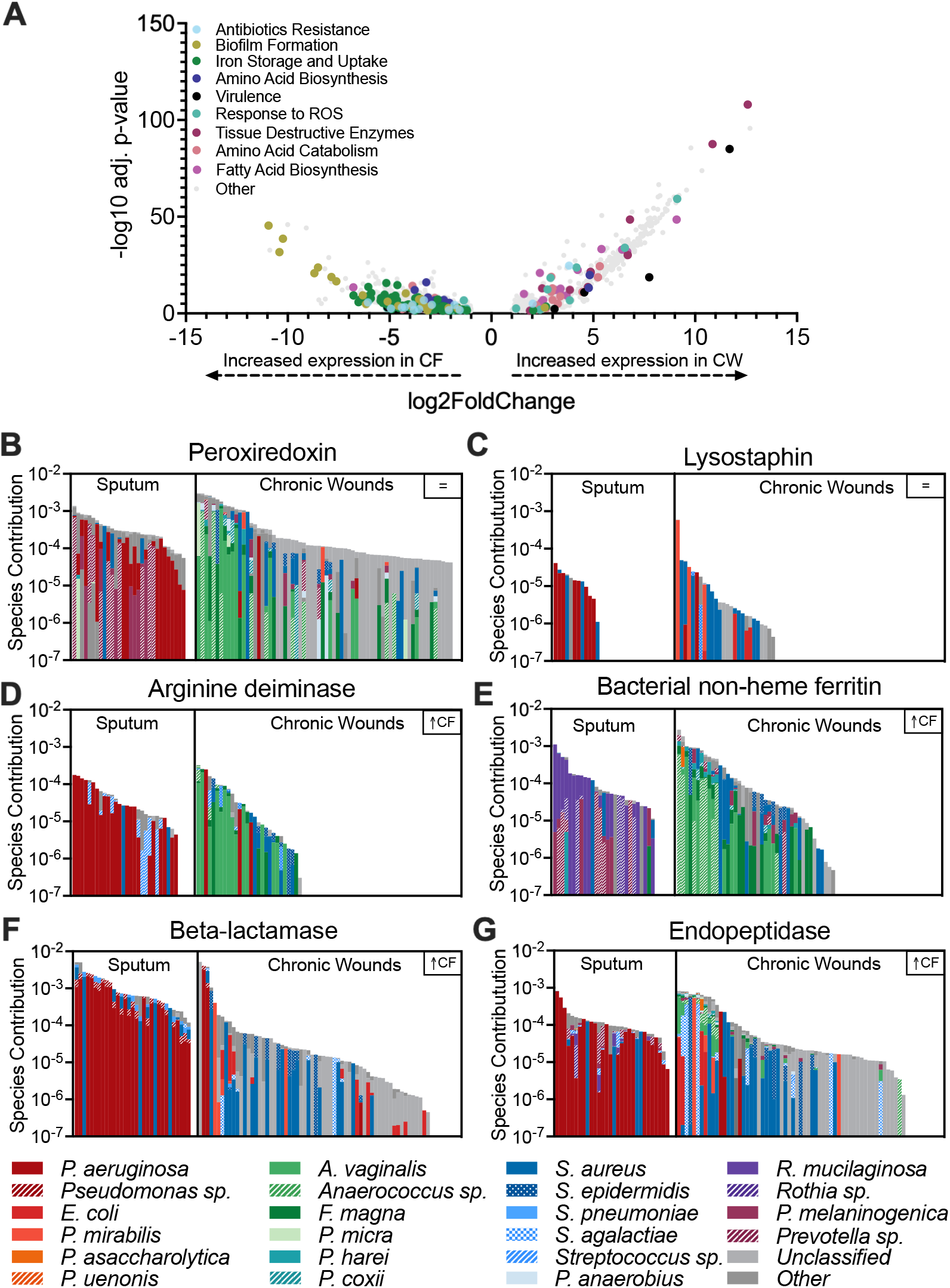
Distinct expression of microbial functions in CF and CW communities. **A)** Volcano plot to highlight differentially expressed functions between infection sites as identified by SAMSA2. 40.37% of the functions were differentially expressed (adjusted *P*-value <0.05, log2FoldChange > 1. **B & C)** Bacterial contribution to the expression of functions conserved across CF and CW environments. **D-G)** Bacterial contribution to the expression of differentially expressed functions.

The increased expression of functions involved in multiple classes of antibiotic resistance in sputum strongly suggests that bacterial community in CF airways may have adapted to negating the effect of the antibiotics used in the management of infection, possibly contributing to the persistence of the lung infection. Further, the enrichment in biosynthetic pathways in CF lung communities indicates these key nutrients are likely limited in this environment. In contrast, the high expression of oxidative stress, tissue destructive enzymes, and catabolic pathways in CW infections indicates the complex community in these infections are degrading host tissue to release nutrients and that nutrients are likely abundant, possibly contributing to bacterial virulence and persistence. This may also be due to the high presence of *S. aureus* in CW infections, which is notorious for synthesizing large quantities of tissue destructive enzymes (16).

## Bacterial community structure and environment influence function

In addition to the distinct functions identified in each infection site, we were interested if the same community members were contributing to conserved functions in each infection, or if distinct community members were contributing to each site. Therefore, we analyzed the stratified output provided by HUMANn3 to evaluate community member contributions. We observed that transcripts were frequently assigned to common pathogens such as *P. aeruginosa, Staphylococcus epidermidis, S. aureus, Streptococcus agalactiae* and anaerobic members of the microbiome such as *Anaerococus vaginalis, Finegoldia magna, Prevotella melaninogenica*, and *Veillonella parvula*. While both groups were prominent contributors to the reduction of oxidative stress and bacterial competition, iron acquisition and biofilm functions were mostly expressed by *P. aeruginosa* in the CF environment while *S. aureus* dominated expression in CW infections. Additionally, tissue degrading enzymes were primarily expressed by *P. aeruginosa* in CF sputum but by the anaerobic microbiota in CW infection. Taken together, our data shows that key community functions are expressed by distinct species in each site, indicating niche differentiation may be occurring during chronic infection. However, it should be noted that one limitation is the short reads used may not allow for species level identification of all functions by HUMANn3.

## Conclusions and Key Takeaways

We found the key functions that drive disease progression in each infection type differ. Further, we showed that the microbial community in each infection type is distinct, and this compositional difference alongside the infection environment is critical in determining functions important for disease progression. Interestingly, we found that the anaerobic microbiota may play a significant role in the progression of chronic infections. Together, these findings will prompt future studies aimed at investigating how co-infecting microbes interact with traditional pathogens, the molecular mechanisms that drive these interactions, and how these interactions impact chronicity.

## Materials and Methods

### Dataset Collection and Validation

We analyzed 102 RNA-sequencing files of chronic wound and cystic fibrosis patients from published studies (7,11,16,17,18,19). We limited our search to metatranscriptomes collected from people with CW in lower extremities & CF and ensured the absence of technical replicates or transcriptomes with reads previously mapped to single bacterial species, which identified 6 studies that fit these criteria. We assessed the quality of the sequence files using FastQC 0.11.9 (20) and removed adapter sequences and reads less than 22 bases with CutAdapt 4.1 (21). Ribosomal RNA sequences were removed with SortMeRNA 4.0.0 (22) using default parameters. The resulting reads were mapped to the human genome (GRCh38/hg38), and processed reads that did not map to GRCh38 were used for community and functional analyses.

### Metatranscriptome Analysis

MetaPhlAn 4.0.1 was used for community composition analysis and to obtain the relative abundances of bacteria in each sample using a minimum read length threshold of 22 bases and other default parameters (23). SAMSA 2.0 and HUMAnN 3.0 were used for functional profiling. First, we analyzed the prokaryotic non-rRNA reads with SAMSA2 to identify the functional profile of the microbial community in each sample (24). SAMSA2 annotated the reads against the RefSeq bacterial database and SEED subsystems database using DIAMOND aligner. Outputs were aggregated and exported for statistical analysis with DESeq2 in RStudio. In addition, we also did functional profiling with HUMAnN3 to obtain the metabolic potential of the microbial communites (25). HUMANn3 uses the DIAMOND aligner to map reads to the UniRef90 database to identify the UniRef protein families, which were regrouped to level 4 enzyme classes (EC). We normalized the reads per kilobase output to relative abundance data with humann_renorm_table and the data was input into MaAsLin2 1.12.0 in RStudio for differential expression analysis.

## Statistical Analyses

All other statistical analyses were performed in RStudio with R version 4.2.2. Data visualizations were performed in GraphPad Prism 9.

## Data Availability

All code used in these analyses is available at https://github.com/Aanuoluwaduro/Metatransriptomics-Microbial-Community-Functions.

The 102 metatranscriptomes used in this study were pulled from the National Center for Biotechnology Information (NCBI) Sequence Read Archive (SRA) under accession numbers: SRP135669, PRJNA573047, PRJNA563930, PRJNA726011, PRJNA576508, PRJNA720438, PRJNA909326.

Additional detailed methods are included in the Supplemental Material.

## Acknowledgments

This study was supported by the National Institutes of Health grant K22AI155927 to C.B.I. Some of the computing for this project was performed at the OU Supercomputing Center for Education & Research (OSCER) at the University of Oklahoma (OU).

## References

1. Peters, B. M., Jabra-Rizk, M. A., O’May, G. A., Costerton, J. W., & Shirtliff, M. E. (2012). Polymicrobial Interactions: Impact on Pathogenesis and Human Disease. Clinical Microbiology Reviews, 25(1), 193–213. https://doi.org/10.1128/CMR.00013-11

2. Ibberson, C. B., & Whiteley, M. (2020). The social life of microbes in chronic infection. Current Opinion in Microbiology, 53, 44–50. https://doi.org/10.1016/j.mib.2020.02.003

3. Murray, J. L., Connell, J. L., Stacy, A., Turner, K. H., & Whiteley, M. (2014). Mechanisms of synergy in polymicrobial infections. Journal of Microbiology, 52(3), 188–199. https://doi.org/10.1007/s12275-014-4067-3

4. Bjarnsholt, T. (2013). The role of bacterial biofilms in chronic infections. APMIS, 121(136), 1–58. https://doi.org/10.1111/apm.12099

5. Burmølle, M., Thomsen, T. R., Fazli, M., Dige, I., Christensen, L., Homøe, P., Tvede, M., Nyvad, B., Tolker-Nielsen, T., Givskov, M., Moser, C., Kirketerp-Møller, K., Johansen, H. K., Høiby, N., Jensen, P. Ø., Sørensen, S. J., & Bjarnsholt, T. (2010). Biofilms in chronic infections – a matter of opportunity – monospecies biofilms in multispecies infections. FEMS Immunology & Medical Microbiology, 59(3), 324–336.

6. Cornforth, D. M., Dees, J. L., Ibberson, C. B., Huse, H. K., Mathiesen, I. H., Kirketerp-Møller, K., Wolcott, R. D., Rumbaugh, K. P., Bjarnsholt, T., & Whiteley, M. (2018). Pseudomonas aeruginosa transcriptome during human infection. Proceedings of the National Academy of Sciences, 115(22), E5125–E5134. https://doi.org/10.1073/pnas.1717525115

7. Pang, M., Zhu, M., Lei, X., Xu, P., & Cheng, B. (2019). Microbiome Imbalances: An Overlooked Potential Mechanism in Chronic Nonhealing Wounds. The International Journal of Lower Extremity Wounds, 18(1), 31–41. https://doi.org/10.1177/1534734619832754

8. Ciofu, O., Hansen, C. R., & Høiby, N. (2013). Respiratory bacterial infections in cystic fibrosis. Current Opinion in Pulmonary Medicine, 19(3), 251. https://doi.org/10.1097/MCP.0b013e32835f1afc

9. Orazi, G., & O’Toole, G. A. (2019). “It Takes a Village”: Mechanisms Underlying Antimicrobial Recalcitrance of Polymicrobial Biofilms. Journal of Bacteriology, 202(1), e00530–19. https://doi.org/10.1128/JB.00530-19

10. Choi, Y., Banerjee, A., McNish, S., Couch, K. S., Torralba, M. G., Lucas, S., Tovchigrechko, A., Madupu, R., Yooseph, S., Nelson, K. E., Shanmugam, V. K., & Chan, A. P. (2019). Co-occurrence of Anaerobes in Human Chronic Wounds. Microbial Ecology, 77(3), 808–820. https://doi.org/10.1007/s00248-018-1231-z

11. Ibberson, C. B., & Whiteley, M. (2019). The Staphylococcus aureus Transcriptome during Cystic Fibrosis Lung Infection. MBio, 10(6), e02774–19. https://doi.org/10.1128/mBio.02774-19

12. Mirković, B., Murray, M. A., Lavelle, G. M., Molloy, K., Azim, A. A., Gunaratnam, C., Healy, F., Slattery, D., McNally, P., Hatch, J., Wolfgang, M., Tunney, M. M., Muhlebach, M. S., Devery, R., Greene, C. M., & McElvaney, N. G. (2015). The Role of Short-Chain Fatty Acids, Produced by Anaerobic Bacteria, in the Cystic Fibrosis Airway. American Journal of Respiratory and Critical Care Medicine, 192(11), 1314–1324. https://doi.org/10.1164/rccm.201505-0943OC

13. Wolcott, R. D., Hanson, J. D., Rees, E. J., Koenig, L. D., Phillips, C. D., Wolcott, R. A., Cox, S. B., & White, J. S. (2016). Analysis of the chronic wound microbiota of 2,963 patients by 16S rDNA pyrosequencing. Wound Repair and Regeneration, 24(1), 163–174. https://doi.org/10.1111/wrr.12370

14. Cuthbertson, L., Walker, A. W., Oliver, A. E., Rogers, G. B., Rivett, D. W., Hampton, T. H., Ashare, A., Elborn, J. S., De Soyza, A., Carroll, M. P., Hoffman, L. R., Lanyon, C., Moskowitz, S. M., O’Toole, G. A., Parkhill, J., Planet, P. J., Teneback, C. C., Tunney, M. M., Zuckerman, J. B., … van der Gast, C. J. (2020). Lung function and microbiota diversity in cystic fibrosis. Microbiome, 8(1), 45. https://doi.org/10.1186/s40168-020-00810-3

15. Lowy, F. D. (1998). Staphylococcus aureus Infections. New England Journal of Medicine, 339(8), 520–532. https://doi.org/10.1056/NEJM199808203390806

16. Fritz, B. G., Kirkegaard, J. B., Nielsen, C. H., Kirketerp-Møller, K., Malone, M., & Bjarnsholt, T. (2022). Transcriptomic fingerprint of bacterial infection in lower extremity ulcers. Apmis, 130(8), 524–534. https://doi.org/10.1111/apm.13234

17. Heravi, F. S., Zakrzewski, M., Vickery, K., Malone, M., & Hu, H. (2020). Metatranscriptomic Analysis Reveals Active Bacterial Communities in Diabetic Foot Infections. Frontiers in Microbiology, 11, 1688. https://doi.org/10.3389/fmicb.2020.01688

18. Lewin, G. R., Kapur, A., Cornforth, D. M., Duncan, R. P., Diggle, F. L., Moustafa, D. A., Harrison, S. A., Skaar, E. P., Chazin, W. J., Goldberg, J. B., Bomberger, J. M., & Whiteley, M. (2023). Application of a quantitative framework to improve the accuracy of a bacterial infection model. Proceedings of the National Academy of Sciences, 120(19), e2221542120. https://doi.org/10.1073/pnas.2221542120

19. Malone, M., Radzieta, M., Peters, T.J., Dickson, H.G., Schwarzer, S., Jensen, S.O., Lavery, L.A.(2021). Host-microbe metatranscriptome reveals differences between acute and chronic infections in diabetes-related foot ulcers. APMIS, 130: 751–762.

20. LaMar, D. (2015). FastQC. https://qubeshub.org/resources/fastqc

21. Martin, M. (2011). Cutadapt removes adapter sequences from high-throughput sequencing reads. EMBnet.Journal, 17(1), Article 1. https://doi.org/10.14806/ej.17.1.200

22. Kopylova, E., Noé, L., & Touzet, H. (2012). SortMeRNA: Fast and accurate filtering of ribosomal RNAs in metatranscriptomic data. Bioinformatics, 28(24), 3211–3217. https://doi.org/10.1093/bioinformatics/bts611

23. Blanco-Miguez, A., Beghini, F., Cumbo, F., McIver, L. J., Thompson, K. N., Zolfo, M., Manghi, P., Dubois, L., Huang, K. D., Thomas, A. M., Piccinno, G., Piperni, E., Punčochář, M., Valles-Colomer, M., Tett, A., Giordano, F., Davies, R., Wolf, J., Berry, S. E., … Segata, N. (2022). Extending and improving metagenomic taxonomic profiling with uncharacterized species with MetaPhlAn 4 (p. 2022.08.22.504593). bioRxiv. https://doi.org/10.1101/2022.08.22.504593

24. Westreich, S. T., Treiber, M. L., Mills, D. A., Korf, I., & Lemay, D. G. (2018). SAMSA2: A standalone metatranscriptome analysis pipeline. BMC Bioinformatics, 19(1), 175. https://doi.org/10.1186/s12859-018-2189-z

25. Beghini, F., McIver, L. J., Blanco-Míguez, A., Dubois, L., Asnicar, F., Maharjan, S., Mailyan, A., Manghi, P., Scholz, M., Thomas, A. M., Valles-Colomer, M., Weingart, G., Zhang, Y., Zolfo, M., Huttenhower, C., Franzosa, E. A., & Segata, N. (n.d.). Integrating taxonomic, functional, and strain-level profiling of diverse microbial communities with bioBakery 3. ELife, 10, e65088. https://doi.org/10.7554/eLife.65088

