## Supplemental Material for "Defining Microbial Community Functions in Chronic Human Infection with Metatranscriptomics"

**Supplementary Material**

**Materials and Methods**

**Dataset Collection:** The datasets used for this study were identified by body part or tissue being affected in samples collected – chronic lower extremity ulcers for CW samples and sputum samples from people with CF, presence of original metatranscriptomic data, presence of comorbidities in patients and treatment course.

**Community Composition Analysis:** We reduced our Metaphlan4 output data to genera and species that had at least one percent abundance prevalence in at least 3 samples.

**Functional Profiling:** Two bioinformatic tools were used for functional profiling. The Simple Annotation of Metatranscriptomes by Sequence Analysis tool **(**SAMSA2) identified 4527 level 4 enzyme classes and the DIAMOND_analsys_counter.py scripts were used to aggregate the outputs which were eventually exported to R studio for statistical analysis. The diversity_stats.R script of the SAMSA2 package was used to compute the mean Shannon and Simpson diversity indices for the two infection communities and the diversity_gaphs.R script was used to plot the graphs of these. We performed differentially expression analysis of the features in the both communities using the run_DESeq_stats.R script. The HMP Unified Metabolic Analysis Network (HUMAnN3) tool, identified 594,273 UniRef protein families which were further regrouped to 2459 unique ECs using the humann_regroup_table command and exported to Rstudio for differential expression analysis.

**Statistical Analyses:**

For the HUMANn3 functional analyses, we performed differential expression analysis using DESeq2 1.38.3 and MaAsLin2 1.12.0 in R studio.

### Figures

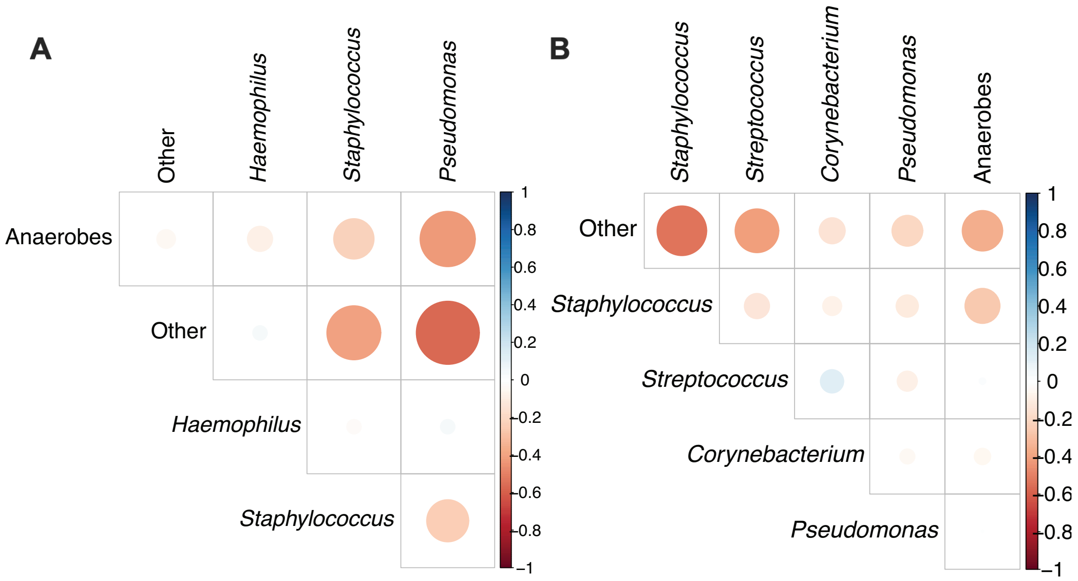


**Figure S1**: Co-occurrence of obligate anaerobes and traditional pathogens in CF and CW samples. **A)** Obligate anaerobes (*Prevotella*, *Actinomyces*, *Veillonella* & *Fusobacterium*) have negative correlation with CF pathogens (*Pseudomonas*, *Staphylococcus*, & *Haemophilus*). **B)** Obligate anaerobes (*Finegoldia, Anaerococcus*, *Peptoniphilus*, *Peptostreptococcus*, *Parvimonas* and *Peptococcus*) have negative correlation with CW pathogens (*Staphylococcus*, *Streptococcus*, *Pseudomonas*, and *Corynebacterium*). **A&B** show the Pearson correlation coefficients between the sum of the relative abundances of the anaerobes and each indicated pathogen. Other bacterial genera present in the community are grouped as other.

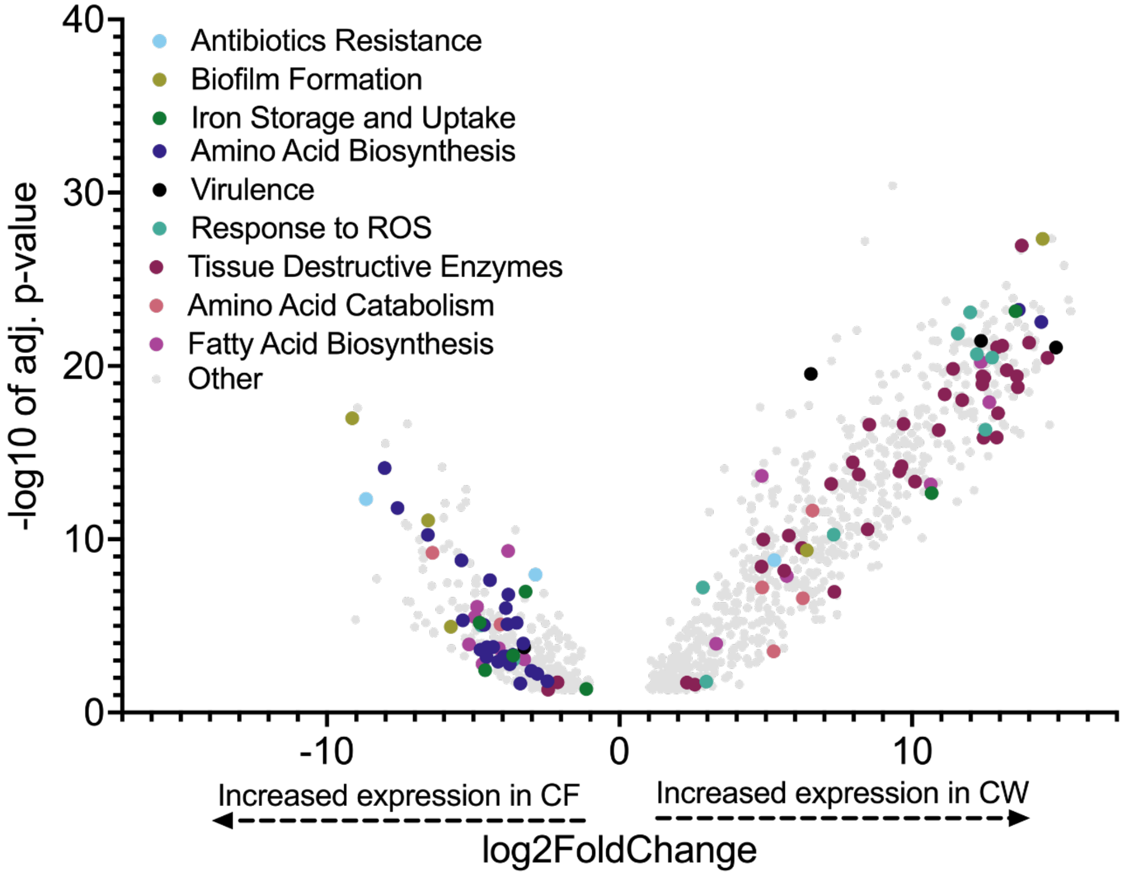


**Figure S2:** Volcano plot to highlight differentially expressed functions in both infection communities as identified by HUMANn3. 42.98% of the functions were differentially expressed (adjusted p-value <0.05, log2FoldChange > 1).

**Supplementary Datasets**

**Dataset S1:** Detailed metadata on all samples used for the study.

**Dataset S2:** Functional analysis data. Sheet 1 - highlighted conserved functions identified with HUMANn3 and SAMSA2 and bacterial contribution to these functions. Sheet 2 - highlighted differentially expressed functions identified with HUMANn3 and SAMSA2 and bacterial contribution to these functions. Sheet 3 - all differentially expressed functions obtained from HUMANn3. Sheet 4 - all differentially expressed functions obtained from SAMSA2.

**Dataset S3**: Pathway analysis data. Sheet 1- all pathway analysis data as identified by HUMANn3. Sheet 2 - all differentially expressed pathways obtained from HUMANn3.
